# The global landscape of SARS-CoV-2 genomes, variants, and haplotypes in 2019nCoVR

**DOI:** 10.1101/2020.08.30.273235

**Authors:** Shuhui Song, Lina Ma, Dong Zou, Dongmei Tian, Cuiping Li, Junwei Zhu, Meili Chen, Anke Wang, Yingke Ma, Mengwei Li, Xufei Teng, Ying Cui, Guangya Duan, Mochen Zhang, Tong Jin, Chengmin Shi, Zhenglin Du, Yadong Zhang, Chuandong Liu, Rujiao Li, Jingyao Zeng, Lili Hao, Shuai Jiang, Hua Chen, Dali Han, Jingfa Xiao, Zhang Zhang, Wenming Zhao, Yongbiao Xue, Yiming Bao

## Abstract

On 22 January 2020, the National Genomics Data Center (NGDC), part of the China National Center for Bioinformation (CNCB), created the 2019 Novel Coronavirus Resource (2019nCoVR), an open-access SARS-CoV-2 information resource. 2019nCoVR features a comprehensive integration of sequence and clinical information for all publicly available SARS-CoV-2 isolates, which are manually curated with value-added annotations and quality evaluated by our in-house automated pipeline. Of particular note, 2019nCoVR performs systematic analyses to generate a dynamic landscape of SARS-CoV-2 genomic variations at a global scale. It provides all identified variants and detailed statistics for each virus isolate, and congregates the quality score, functional annotation, and population frequency for each variant. It also generates visualization of the spatiotemporal change for each variant and yields historical viral haplotype network maps for the course of the outbreak from all complete and high-quality genomes. Moreover, 2019nCoVR provides a full collection of SARS-CoV-2 relevant literature on COVID-19 (Coronavirus Disease 2019), including published papers from PubMed as well as preprints from services such as bioRxiv and medRxiv through Europe PMC. Furthermore, by linking with relevant databases in CNCB-NGDC, 2019nCoVR offers data submission services for raw sequence reads and assembled genomes, and data sharing with National Center for Biotechnology Information. Collectively, all SARS-CoV-2 genome sequences, variants, haplotypes and literature are updated daily to provide timely information, making 2019nCoVR a valuable resource for the global research community. 2019nCoVR is accessible at https://bigd.big.ac.cn/ncov/.

## Introduction

The severe respiratory disease COVID-19 [1], since its outbreak in late December 2019, has rapidly spread as a pandemic. As of 14^th^ July 2020, 12,964,809 confirmed cases have been reported in 216 countries/territories/areas (WHO Situation Report Number 176; https://www.who.int/emergencies/diseases/novel-coronavirus-2019/situation-reports/). As the causative agent of COVID-19, SARS-CoV-2 samples have been extensively isolated and sequenced by different countries and laboratories [2], resulting in a considerable number of viral genome sequences worldwide. Therefore, public sharing and free access to a comprehensive collection of SARS-CoV-2 genome sequences is of great significance for worldwide researchers to accelerate scientific research and knowledge discovery and also help develop medical countermeasures and sensible decision-making [3].

To date, unfortunately, SARS-CoV-2 genome sequences generated worldwide were scattered around different database resources, primarily including the Global Initiative on Sharing All Influenza Data (GISIAD) [4] repository and NCBI GenBank [5]. Many sequences exist in multiple repositories but their updates are not synchronized. This makes it extremely challenging for worldwide users to effectively retrieve a non-redundant and most updated set of sequence data, and to collaboratively and rapidly deal with this global pandemic. Towards this end, we constructed the 2019 novel coronavirus resource (2019nCoVR, https://bigd.big.ac.cn/ncov/) in CNCB-NGDC, with the purpose to provide public, free, rapid access to a complete collection of non-redundant global SARS-CoV-2 genomes by comprehensive integration and value-added annotation and analysis [6]. Since its inception on 22 January 2020, 2019nCoVR is updated on daily basis, leading to unprecedentedly dramatic data expansion from 86 genomes in its first release to 64,789 genomes in its current version (as of 14^th^ July 2020). Moreover, it has been greatly upgraded by equipping with enhanced data curation and analysis pipelines and online functionalities, including data quality evaluation, variant calling, variant spatiotemporal dynamic tracking, viral haplotype construction, and interactive visualization with more friendly web interfaces (Table 1). Here we report these significant updates of 2019nCoVR and present the global landscape of SARS-CoV-2 genomes, variants and haplotypes.

**Table 1.** Comparison of functional modules between two versions of 2019nCoVR

| Functionality | Version 1 | Version 2 |
| --- | --- | --- |
| Integration of coronavirus sequences | ✓ | ✓ |
| Genomic sequences and metadata of 2019-nCoV | ✓ | ✓ |
| Variant statistics and visualization of 2019-nCoV | ✓ | ✓ |
| Phylogenetic tree | ✓ | ✓ |
| Raw sequence data submission | ✓ | ✓ |
| Genome assembly submission | ✓ | ✓ |
| Literature |  | ✓ |
| Clinical information |  | ✓ |
| Sequence integrity and quality assessment |  | ✓ |
| Variant annotation based on 3D structures (S Protein) |  | ✓ |
| Spatiotemporal dynamics analysis of genomic variants |  | ✓ |
| Haplotype network and dynamic evolution |  | ✓ |
| AI diagnosis and online tools |  | ✓ |
| Enhanced user-friendly interface |  | ✓ |

### Database content and features

#### Statistics of SARS-CoV-2 genome assemblies

Since the outbreak of COVID-19, the number of SARS-CoV-2 genome sequences released globally has been increasing at an unprecedented rate. To facilitate public free access to all genome assemblies and help worldwide researchers better understand the variation and transmission of SARS-CoV-2, we perform daily updates for 2019nCoVR by integrating all available genomes throughout the world and conducting value-added curation and analysis (**Figure 1**). As of 14^th^ July 2020, 2019nCoVR hosted a total of 64,789 non-redundant genome sequences and provided a global distribution of SARS-CoV-2 genome sequences in 97 countries/regions across 6 continents (Figure 1A). Duplicated sequences from different databases are merged with all IDs cross-referenced. Sequences are contributed primarily by United Kingdom (28,823, 44.5%), United States (13,556, 20.9%), Australia (2351, 3.6%), Spain (1852, 2.9%), Netherlands (1605, 2.5%), India (1581, 2.4%), and China (1431, 2.2%). According to our statistics, SARS-CoV-2 genome sequences started to grow rapidly from mid-March (https://bigd.big.ac.cn/ncov/release_genome), concordant with the outset of global pandemic of COVID-19. A full list of our sequence dataset including strain name, accession number and source is provided in Table S1.

**Figure 1.**
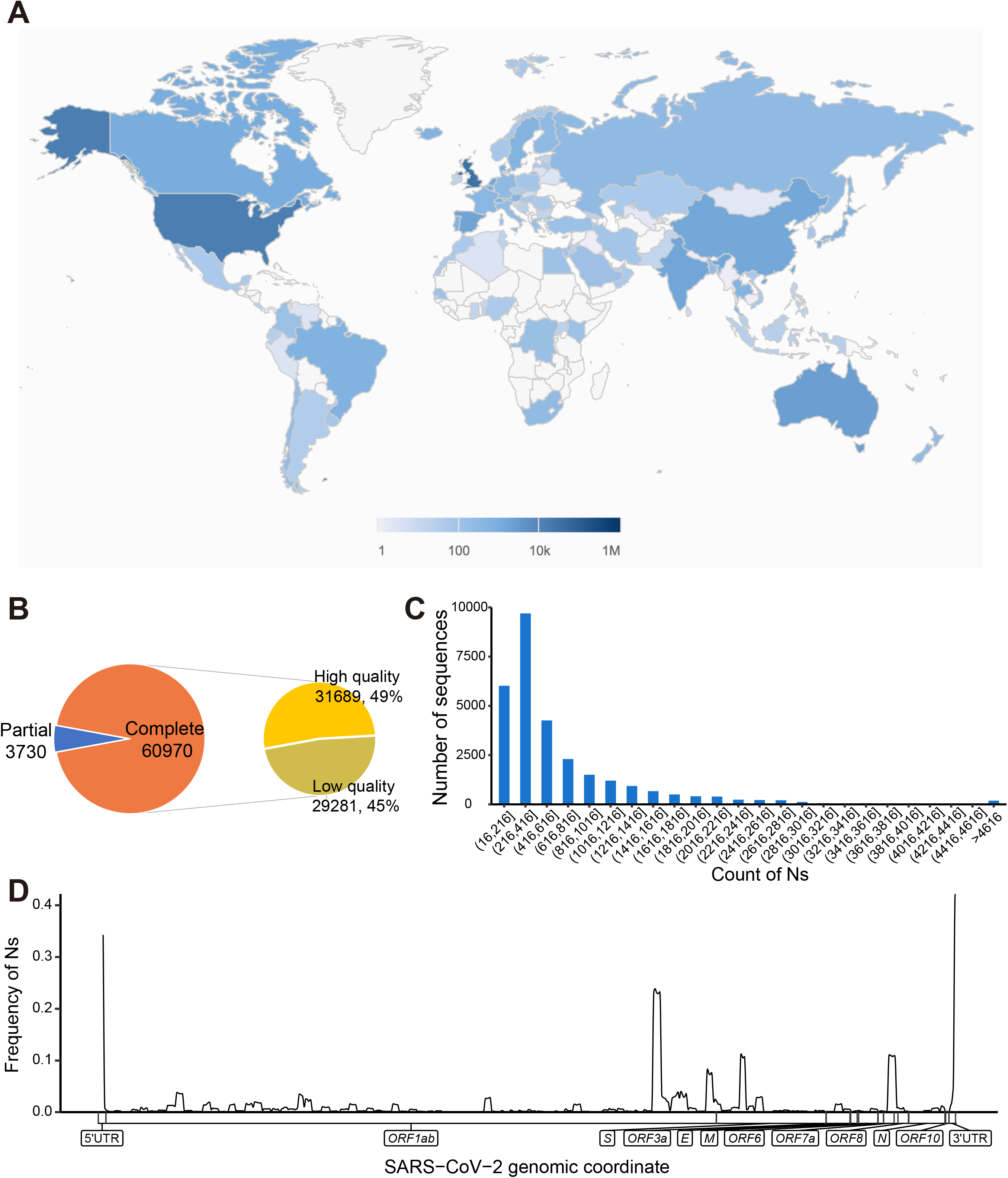
Statistics and distribution of all released SARS-CoV-2 genomes. (A) Distribution of released genome sequences by country, territory or region; (B) Number and percentage of complete and high-quality genomes; (C) Sequences number of different Ns ranges for low-quality genomes; (D) Frequency distribution of Ns across the whole genome.

To provide high-quality genome sequences that are critically essential for downstream analyses (ranging from variant calling to haplotype construction), we perform sequence integrity and quality assessment for all newly collected sequences. Among all released human-derived genome sequences (64,700), 60,970 (94.2%) are complete, and 31,689 (49%) are high-quality (with high coverage) (Figure 1B). Most of the low-quality sequences (29,281, 99.7%) contain different numbers of unknown bases (Ns). Among these sequences, 60% have 16-500 Ns (median 258), and 40% have more than 500 Ns (Figure 1C). Further investigation of the genomic locations reveals that some genomic regions have high coverage of Ns (Figure 1D). Sequence integrity and quality assessment results are available for all genome sequences, and can be used as filters for sequence browse and search.

#### Landscape of genomic variants

Bases on 31,685 globally human-derived high-quality complete genome sequences (in what follows, only high-quality complete genome sequences are used for downstream analysis if not indicated otherwise), we investigate the landscape of SARS-CoV-2 genomic variants by comparison with the reference genome (MN908947.3 in NCBI) (**Figure 2**). By 14^th^ July 2020, a total of 13,428 variants were identified, including 12,828 (95.5%) single nucleotide polymorphisms (SNPs), 437 deletions, 116 insertions, and 47 indels (a combination of an insertion and a deletion, affecting 2 or more nucleotides) (Figure 2A). More than half of these SNPs (6770, 50.4%) are nonsynonymous, causing amino acid changes. To gain the functional effects of those missense variants of S spike protein from the perspective of spatial location (e.g. key functional domain or binding region), mutated amino acids are projected onto protein 3D structures, which can be viewed by 360 degree rotation (Figure 2B). We further explore the distributions of these variants across different genes. Noticeably, the three genes *ORF1ab, S*, and *N* accumulate more variants (Figure 2C) and SNP densities (i.e., the number of mutations per nucleotides in the gene region) are higher in several gene regions including *ORF7a, ORF3a, ORF6* and *N*(https://bigd.big.ac.cn/ncov/variation/annotation).

**Figure 2.**
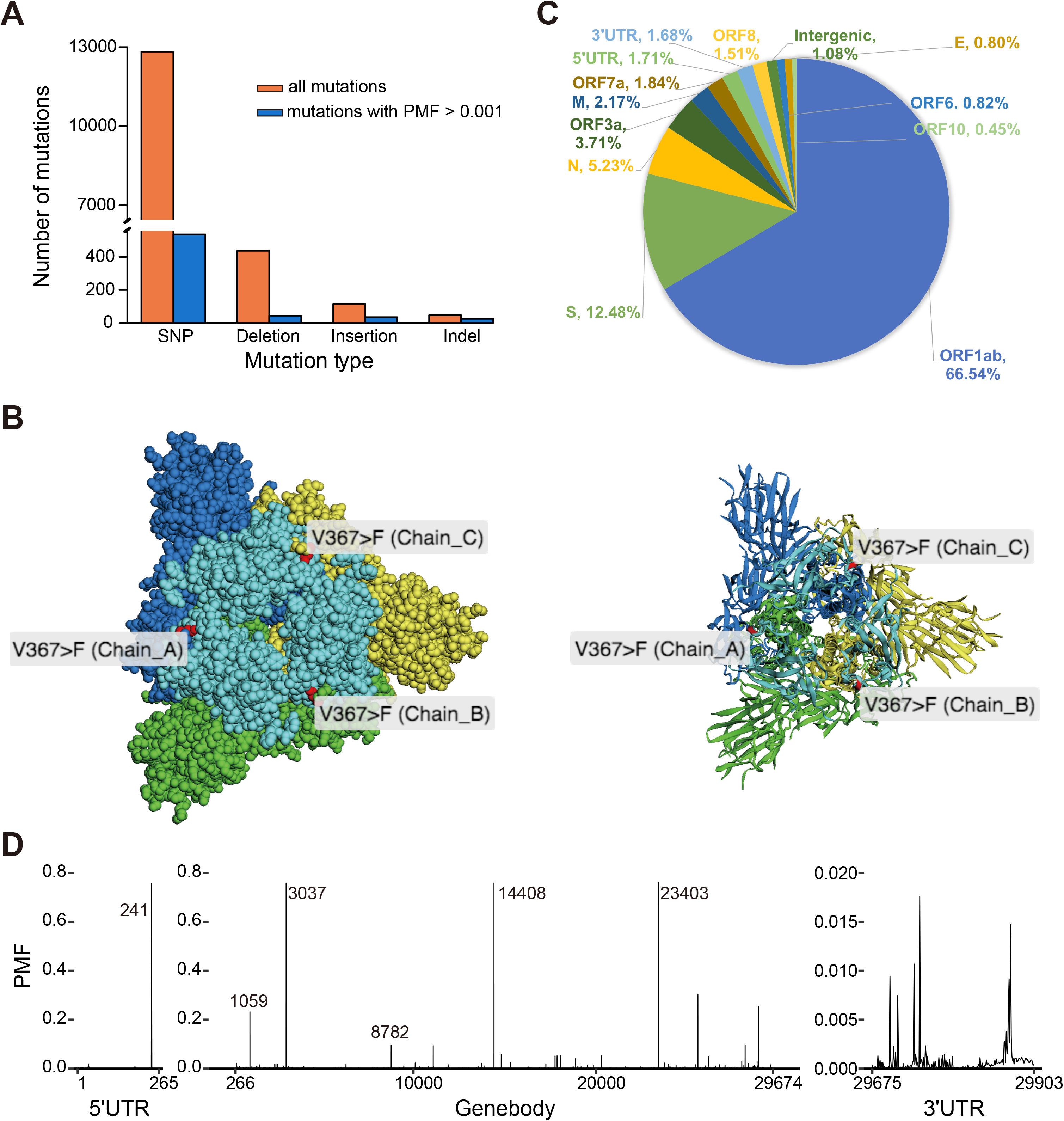
Landscape of genomic variants. (A) Numbers of different mutation types (including SNPs, deletion, insertion, and indels; the orange bar represents all mutations while the blue bar represents mutations with PMF>0.001); (B) Structure display for nonsynonymous mutations; (C) Pie chart of variant annotation for each gene; (D) Population mutated frequency (PMF) for all variants.

For each variant, we investigate its population mutation frequency (PMF, the ratio of the number of mutated genomes to the total number of complete high-quality genomes) (Figure 2D). Clearly, there are 62 variants with PMF > 1%, 18 variants with PMF > 5%, and 4 variants with PMF > 75.8% (that is, position 241 in 5’UTR, positions 3037 and 14,408 in *ORF1ab*, and 23,403 in *S*), potentially representing main prevalent virus genotypes across the global. All identified variants and their functional annotations are publicly accessible and an online pipeline for variant identification and functional annotation is provided and freely available at https://bigd.big.ac.cn/ncov/analysis.

#### Spatiotemporal dynamics of genomic variants

To track the dynamics of SARS-CoV-2 genomic variants, particularly *de novo* mutations, we explore the spatiotemporal change of population frequency for each variant according to sampling time and location (**Figure 3**). Among the 18 sites with PMF > 5%, some are mutated simultaneously and in a linkage manner (Figure 3A), such as mutations at positions 8782 and 28,144 reported in [7]. Specifically, these two sites appeared in the early stage of the outbreak since 30 December 2019, and their mutation frequencies reach ~33% around 22 January 2020, and then gradually decline to 9.6% currently. Contrastingly, some variants appear only since the middle stage around 3 March 2020; such as the mutation at position 23,403 (provoking an amino acid change D614G of the *S* protein), is accompanied by three other mutations, namely, a C-to-U mutation at position 241 in the 5’UTR, a silent C-to-U mutation in the gene *nsp3* at position 3037, and a missense C-to-U mutation in the gene *RdRp* at position 14,408 (P4715L). To facilitate users to investigate any variant of interest, we provide an interactive heatmap in 2019nCoVR (https://bigd.big.ac.cn/ncov/variation/heatmap) to dynamically display and cluster the mutation patterns over all sampling dates, with customized options available that allow users to select specific variant frequency, annotated gene/region, variant effect type, and transcription regulation sequence (TRS).

**Figure 3.**
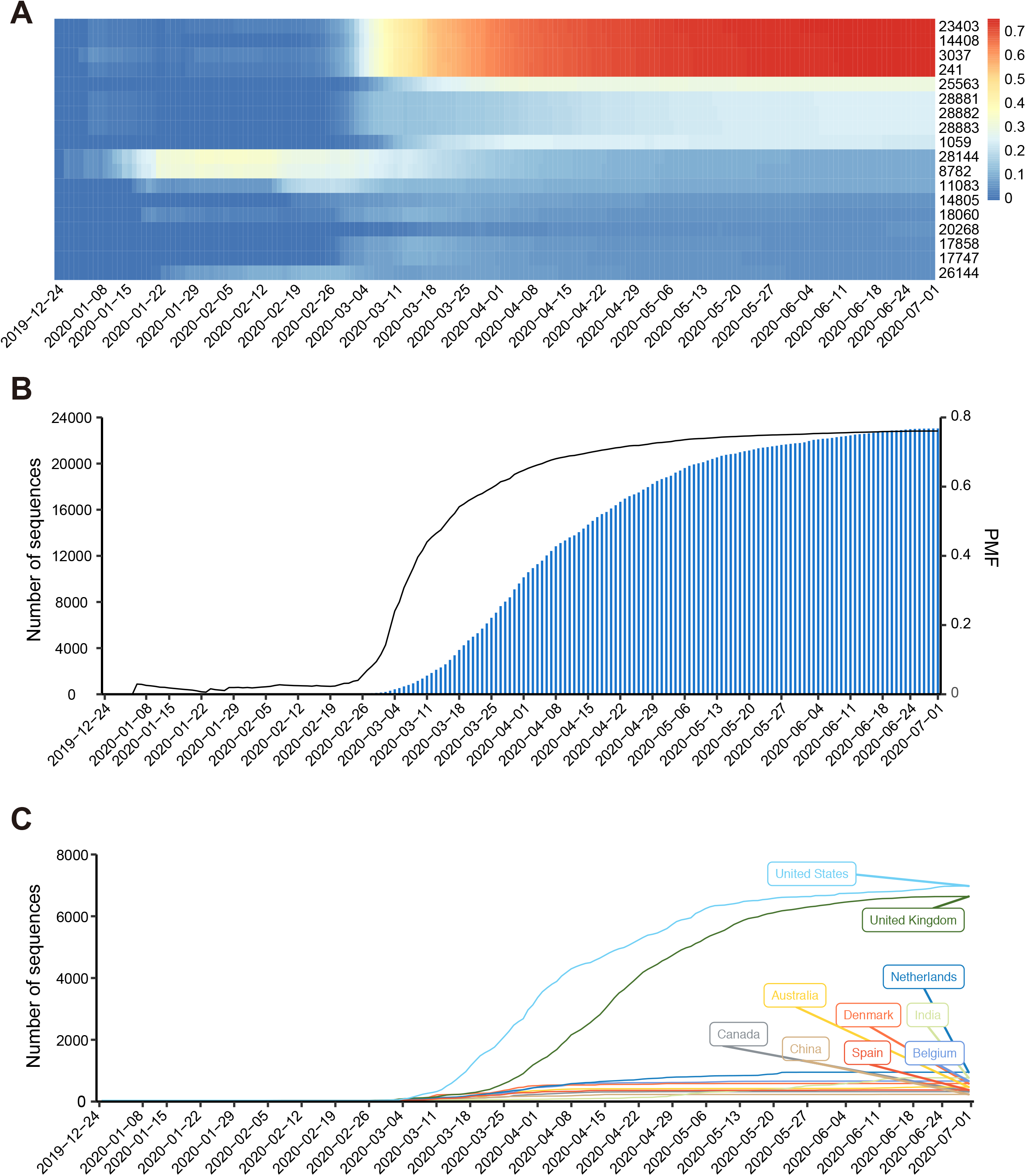
Spatiotemporal dynamics of genomic variants. (A) Population mutation frequency (PMF) of variants over time; (B) PMF and cumulative sequence growth curve for position (n23403, pD614G); (C) Cumulative growth curve of the number of mutated sequences in selected countries for position (n23403, pD614G).

Moreover, we investigate dynamic patterns of SARS-CoV-2 genomic variants across different sampling locations over time. Taking the variant at position 23,403 (D614G) as an example, its PMF has dramatically increased from 0 at the end of February to 76.2% right now, and the mutation pattern G614 has been gradually dominant along with the development of pandemic (Figure 3B), presumably indicating that the mutated genotypes may have higher transmissibility[8]. In terms of the absolute number of mutation patterns across different countries/regions, G614 emerges dominantly in Europe and North America (Figure 3C). (https://bigd.big.ac.cn/ncov/variation/annotation/variant/23403). When investigating the mutation pattern for each country (**Figure 4**), we find that sequences from some Asian countries (such as South Korea, Malaysia, and Nepal) have no or very few G614 mutation, whereas those from Europe and America (e.g. Argentina, Czech Republic and Serbia) do have the G614 pattern that is dominated among contemporary samples. In some countries, both the D614 and G614 patterns are co-circulating early in the epidemic, but the mutated pattern soon begins to be dominant such as in Australia, Belgium, Canada, Chile, France, Israel, United States and United Kingdom [8]. The accumulation of this mutation varies in different parts of the world, possibly due to the prevention and control measures adopted by some countries/regions. Taken together, 2019nCoVR features spatiotemporal dynamics tracking of SARS-CoV-2 genomic variants and thus bears great potential to help decipher viral transmission and adaptation to the host [8].

**Figure 4.**
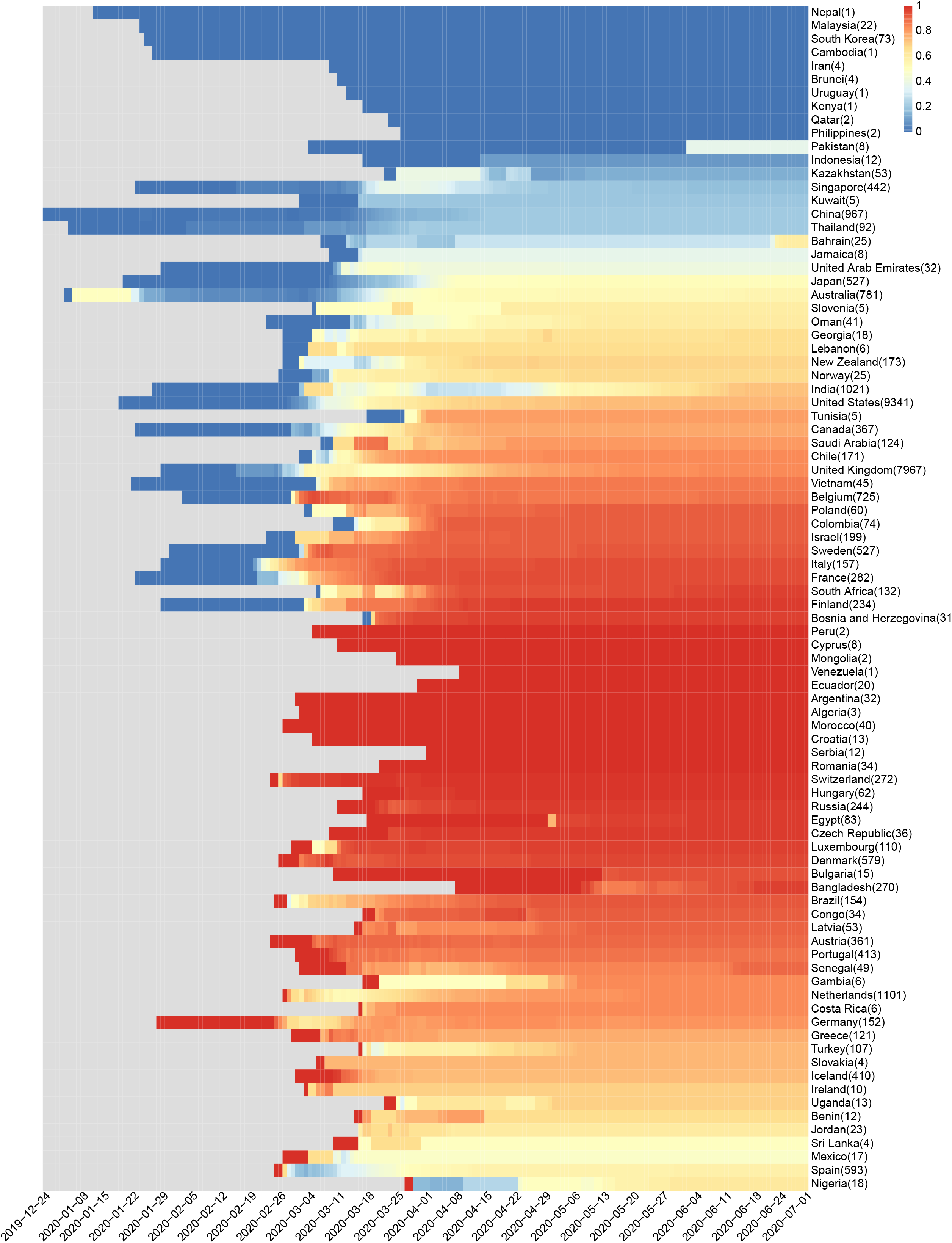
The population mutated frequency (PMF) of G614 for each country over time.

#### Haplotype network construction and characterization

To better characterize the diversity of virus sequences, we build SARS-CoV-2 haplotypes based on all identified variants beyond UTRs regions. As a result, 17,624 haplotypes were identified from 31,685 complete high-quality genome sequences as of 14^th^ July 2020. Based on this, we construct a haplotype network for SARS-CoV-2 (**Figure 5**), a graphical representation of genomic variations by inferring relationships between individual genotypes, according to the principle of the shortest set of connections that link all nodes (genotypes) where the length of each connection represents the genetic distance [9]. To provide a whole picture of the pandemic transmission in a spatiotemporal manner, we visualize the SARS-CoV-2 haplotype network by sample collection date and across different countries/regions. It not only allows users to easily obtain a landscape of SARS-CoV-2 haplotypes and their relationships, but also helps users to navigate a set of haplotypes for a specific country/region linking with additional associated information such as the number of genomes, sampling time and location (Figure 5A).

**Figure 5.**
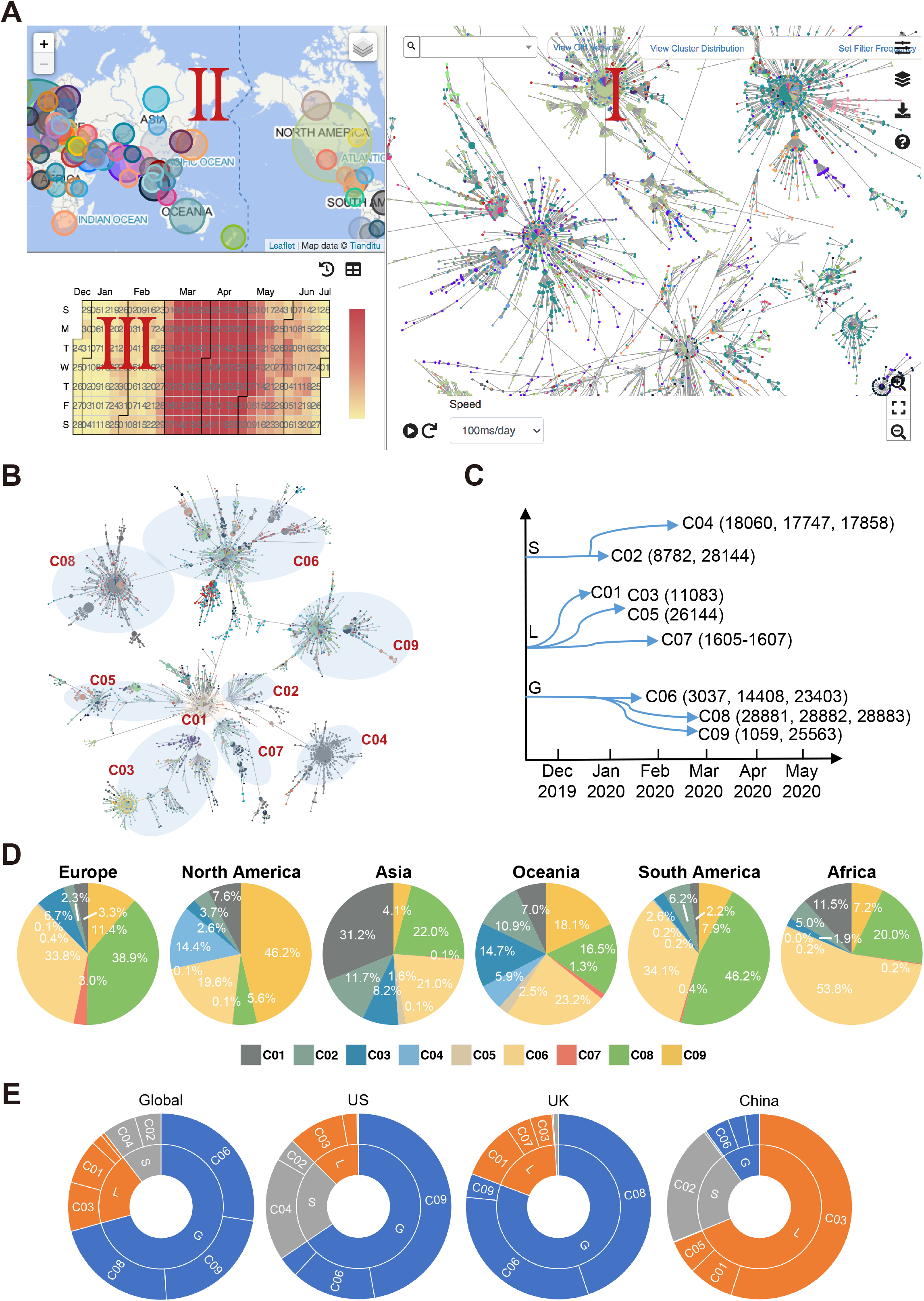
Haplotype network and cluster identification and distribution. (A) The snapshot of haplotype network dashboard, which can dynamically show the development of haplotype (I) across countries (II) and over time (III). Each node in the network represents a haplotype and the node size is proportional to the number of viral genome sequences, where the edge between any two nodes represents the genetic distance between two haplotypes (i.e. the number of mutation sites); (B-C) Schematic diagram of haplotype clusters (C01–C09) and their corresponding common mutation sites for each cluster; (D) Distribution of C01–C09 clusters across different continents; (E) Distribution of different clusters throughout the world and in three representative countries (US, UK and China).

According to the haplotype network, we classify all genome sequences into nine major clusters (labelled as C01–C09; see Methods for details) (Figure 5B, 5C; **Table 2**). As the ongoing pandemic spread of SARS-CoV-2, new branches that evolve and spread faster are gradually emerging, such as clusters C04, C06, C08, and C09 (Table 2). The dominant clusters are C06 (8681, 27.4%), C08 (7,889, 24.9%), and C09 (6,940, 21.9%) (Figure 5D), which are characterized by those signature mutations of C241T, C3037T, C14408T, and A23403G, and are defined as the G clade (as the mutation at position 23,403 provoking an amino acid change D614G of S protein). These sequences have spread to 82 countries worldwide, and become the main epidemic virus type in most countries in Europe, North America, South America, Africa and West Asia, etc. For example, there were about 6827 (71.5%), 8305 (83.4%), and 970 (18.5%) sequences originated from the G clade in the United States, United Kingdom, and China, respectively (Figure 5E). The wide spread and prevalence of this clade in different countries may suggest the adaptability of the virus type to human [8].

**Table 2.** Signature mutations of haplotype clusters

| Clade | Cluster ID | Genomic Location | Gene/Genomic Regions | Mutation | Amino acid Position and Change | Mutation Frequency | Number of Mutated Sequences | Classification in Lu et.al 2020 NSR<sup>a</sup> | Lineage in Rambaut et.al 2020 Nat Microbiol<sup>b</sup> |
| --- | --- | --- | --- | --- | --- | --- | --- | --- | --- |
| L | C01 | NA | NA | NA | NA | NA | NA | L | B/B.3/B.1.3 etc. |
|  | C03 | 11083 | *ORF1ab* | G->T | p.3606L>F | 0.09 | 2982 | L | B.2/B.2.1/B.4 etc. |
|  | C05 | 26144 | *ORF3a* | G->T | p.251G>V | 0.05 | 1592 | L | B/B.2 |
|  | C07 | 1604 | *ORF1ab* | AATG->A | p.447-448ND>N | 0.02 | 503 | L | B/B.8 |
| S | C02 | 8782 | *ORF1ab* | C->T | p.2839S | 0.09 | 3034 | S | A/A.3/A.4/A.5 |
|  |  | 28144 | *ORF8* | T->C | p.84L>S | 0.09 | 3063 |  |  |
|  | C04 | 17747 | *ORF1ab* | C->T | p.5828P>L | 0.05 | 1644 | S | A.1/A.1.1/A.1.2 |
|  |  | 17858 | *ORF1ab* | A->G | p.5865Y>C | 0.05 | 1657 |  |  |
|  |  | 18060 | *ORF1ab* | C->T | p.5932L | 0.05 | 1695 |  |  |
| G | C06 | 241 | *5'UTR* | C->T |  | 0.75 | 24028 | L | B.1/B.1.5/B.1.11 etc. |
|  |  | 3037 | *ORF1ab/nsp4* | C->T | p.924F | 0.75 | 24045 |  |  |
|  |  | 14408 | *ORF1ab/RdRp* | C->T | p.4715P>L | 0.75 | 24055 |  |  |
|  |  | 23403 | *S* | A->G | p.614D>G | 0.76 | 24128 |  |  |
|  | C08 | 28881 | *N* | G->A | p.203R>K | 0.25 | 8003 | L | B.1/B.1.1/B.1.10 etc. |
|  |  | 28883 | *N* | G->C | p.204G>R | 0.25 | 7995 |  |  |
|  |  | 28882 | *N* | G->A | p.203R | 0.25 | 7985 |  |  |
|  | C09 | 1059 | *ORF1ab* | C->T | p.265T>I | 0.23 | 7357 | L | B.1/B.1.21/B.1.43 |
|  |  | 25563 | *ORF3a* | G->T | p.57Q>H | 0.30 | 9594 |  |  |

Note: <sup>a</sup> Tang X, Wu C, Li X, Song Y, Yao X, Wu X, et al. On the origin and continuing evolution of SARS-CoV-2. National Science Review 2020.
<sup>b</sup> Rambaut A, Holmes EC, O'Toole A, Hill V, McCrone JT, Ruis C, et al. A dynamic nomenclature proposal for SARS-CoV-2 lineages to assist genomic epidemiology. Nat Microbiol 2020.

#### Implementation

2019nCoVR was built based on B/S (Browser/Server) architecture. In the browser-side, it was developed by JSP (Java Server Pages), HTML, CSS, AJAX (Asynchronous JavaScript and XML), JQuery (a cross-platform and feature-rich JavaScript library; http://jquery.com) as well as Semantic-UI (an open source web development framework; https://semantic-ui.com). In the server-side, it was implemented by using Spring Boot (a rapid application development framework based on Spring; https://spring.io). For data storage, MySQL (https://mysql.com) was used. For interactive visualization, HighCharts (a modern SVG-based, multi-platform charting library; https://highcharts.com), D3.js (a JavaScript library for manipulating documents based on data; https://d3js.org) and 3Dmol.js (a JavaScript library for visualizing protein structure associated with mutated amino acids) [21] were employed in 2019nCoVR. The haplotype network was implemented by d3js, Leaflet (http://leafletjs.com), and Echarts (http://echarts.baidu.com/).

## Discussion

Genome sequencing is vital to understand the epidemiology of SARS-CoV-2, since it is not only useful for deciphering its genome sequences and investigating its evolution and transmission, but also highly effective at determining whether individuals are part of the same transmission chain [10]. According to 2019nCoVR, however, the ratio of sequenced samples to the number of confirmed cases is very low in some countries/regions (Figure S1), and even genome sequences are unavailable in some affected countries/regions. The SARS-CoV-2 sampling bias and depth may lead to inaccurate transmission patterns and phylogenetic relationships [11]. Based on sequencing all infected cases in a single region, it has proved that the transmission of *Clostridium difficile* from symptomatic patients accounts for only one third of all infected cases [12]. As our current understanding is still very limited, we call for more efforts and collaborations in sequencing more SARS-CoV-2 genomes from both symptomatic and asymptomatic cases.

Besides, as these released SARS-CoV-2 genome sequences were generated by multiple different laboratories on different sequencing platforms, the quality of genome sequences is another important factor, such as the Ns of genome, which may affect variant calling and biased population frequency estimation. As mentioned in results, the frequency of Ns in some genomic regions is high, possibly due to the low sequencing coverage, low-complexity of sequence, low-efficiency of PCR primers used in sequencing library construction, secondary structure of RNA, etc. However, most of the sequencing coverage information is unavailable, making it challenging to evaluate whether the Ns were due to low sequencing coverage. By further investigating those genomic regions with high frequency of Ns, we found (1) their GC and AG contents are close to the average of the whole genome, excluding the possibility of low complexity of sequence; (2) the length of these regions ranges from 210 to 320 bp (similar to the length of PCR product) and more than 60% of the related sequences are generated on Illumina platform (based on PCR amplification), suggesting that those Ns regions may be resulted from low-efficiency of PCR Primers during sequencing library construction; (3) by analyzing the secondary structure of these regions’ RNA sequences, we found the minimum free energy is lower than those randomly extracted regions, indicating that the secondary structure is more stable and may affect the determination of genome sequences(Figure S2). Our future efforts are to construct a recognized benchmark for quality assessment and data filtration.

Compared to the early overly simplified L-S classification [8] and those comprehensive lineages defined by Rambaut et al. [13], our classification scheme with nine clusters provides a moderate system that can be correlated with the others (Table 2). The nine clusters could also be grouped into three clades defined in [8, 10], namely, S (C02 and C04), G (C06, C08 and C09), and L (the rest clusters). Although haplotype network cannot give a precise evolutionary position as phylogenetic trees do, it can be used to quickly inform the clustering of viruses according to signature mutations in each haplotype. Definitely, new clusters will be introduced as the virus is continuing to evolve.

A data-driven response to SARS-CoV-2 requires a public, free, and open-access data resource that contains complete high-quality genome sequence data, and equips with automated online pipelines to rapidly analyze genome sequences. Thus, 2019nCoVR (together with other resources in CNCB-NGDC) provides a wide range of data services, involving raw sequencing data archive, genome sequence and meta information management with quality control and curation, variation analysis and data presentation and visualization. Additionally, to facilitate worldwide users to monitor any variant that may be associated with rapid transmission and high virulence, 2019nCoVR R, when compared to GISAID and NCBI Virus, features spatiotemporal dynamic tracking for all identified variants. To better understand the epidemiology of SARS-CoV-2, future directions are to collect ever more genome sequences worldwide, include other types of omics data (such as transcriptome and epitranscriptome, if available) [14] and also provide more friendly interfaces and online tools in support of worldwide research activities.

## Methods

### Data collection and integration

All genome sequences as well as their related metadata were integrated from SARS-CoV-2 resources worldwide, including NCBI [5], GISAID [15], CNCB-NGDC [16], NMDC [17] and CNGB [18]. To provide a non-redundant dataset, duplicated records from different databases were identified and merged.

### Quality control and curation

To determine the integrity of genome sequences, one sequence is defined as ‘Complete’ if it is longer than 29000 bases and covers all protein-coding/CDS regions of SARS-CoV-2 (bases 266:29674 of GenBank: MN908947.3); otherwise, it is defined as “Partial”. Furthermore, to examine the quality of genome sequences, unknown bases (Ns) and degenerate bases (Ds, more than one possible base at a particular position and sometimes referred as “mixed bases”) were counted for each sequence. By our default definition, one sequence is “high-quality” if Ns≦15 and Ds≦50, and “low-quality” otherwise. Besides, any sequence is clearly labelled when the number of variants≧15 or the total number of deletion≧2 or the distribution of sequence variation is more aggregated (the ratio of the number of variants divided by the total number of bases in a window≧0.25).

### Variant identification and haplotype network construction

Only complete and high-quality genome sequences were used for downstream analyses, including sequence comparison, variant identification, functional annotation, and haplotype network construction. Genome sequence alignment was performed with Muscle (3.8.31) [19] by comparing against the earliest released SARS-CoV-2 genome (MN908947.3). Sequence variation was identified directly using an in-house Perl program. The effect of variants was determined using VEP (ENSEMBL Variant Effect Predictor) [20].

SARS-CoV-2 haplotypes were constructed based on short pseudo sequences that consist of all variants (filtering out variations located in UTR regions) only. Then, all these pseudo sequences were clustered into groups, and each group (a haplotype) represents a unique sequence pattern. The haplotype network was inferred from all identified haplotypes, where the reference sequence haplotype was set as the starting node, and its relationship with other haplotypes was determined according to the inheritance of mutations. As a result, nine major haplotype network clusters (denoted as C01–C09) were obtained according to the phylogenetic tree-and-branch structure and those shared landmark mutations (Table 1). Specifically, mutations with PMF≧5% (except for ATG deletion at position1605, PMF≈3%) were selected, and those co-occurred mutations were determined by LD linkage analysis. A cluster was referred to sequences with those co-occurred landmark mutations.

### Data availability

SARS-CoV-2 genomes, variants (in vcf format) and their annotations are publicly available at https://bigd.big.ac.cn/ncov/.

### CRediT author statement

**Shuhui Song:** Conceptualization, Methodology, Data Analysis, Writing Original Paper, Reviewing and Editing **Lina Ma:** Data curation, Methodology, Writing **Dong Zou:** System Development, Writing **Dongmei Tian:** Methodology, Data Analysis **Cuiping Li:** Methodology, Data Analysis **Junwei Zhu:** System Development **MeiliChen:** Data curation **Anke Wang**: System Development **Yingke Ma:** System Development **MengWei Li:** Methodology, System Development **Xufei Teng:** Visualization **Ying Cui:** Data curation **Guangya Duan:** Data curation **Mochen Zhang:** Data curation **Tong Jin:** Data curation **Chengmin Shi:** Methodology **Zhenglin Du:** Methodology **Yadong Zhang:** Methodology **Chuandong Liu:** Methodology **Rujiao Li:** Data curation **Jingyao Zeng:** Data curation **Lili Hao:** Data curation **Shuai Jiang:** Methodology **Hua Chen:** Supervision **Dali Han:** Supervision **Jingfa Xiao:** Supervision, Methodology **Zhang Zhang:** Conceptualization, Supervision, Reviewing and Editing **Wenming Zhao:** Conceptualization, Supervision, Methodology **Yongbiao Xue:** Conceptualization, Supervision **Yimin Bao:** Conceptualization, Supervision, Reviewing and Editing

## Acknowledgements

We thank our colleagues and students for their hard working on the 2019nCoVR (https://bigd.big.ac.cn/ncov). We also thank a number of users and CNCB-NGDC members for reporting bugs and sending comments. Complete genome sequences used for analyses were obtained from the CNCB-NGDC, CNGBdb, GenBank, GISAID, and NMDC databases. We acknowledge the sample providers and data submitters listed on Table S1

## Supplementary Figure and Table legend

Figure S1 Distribution of genome sequence count divided by the number of confirmed cases for each country.

Figure S2 Compositional analysis of whole genome and two representative genomic regions with high frequency of Ns. (A) GC and AG Compositional variability; (B) Two representative genomic regions with high frequency of Ns and distribution of sequencing platforms for those corresponding sequences; (C) The secondary structure of one representative Ns region and one non-Ns region.

Table. S1 Coronavirus sequence datasets used for the study.

## References

[1] Coronaviridae Study Group of the International Committee on Taxonomy of Viruses. The species Severe acute respiratory syndrome-related coronavirus: classifying 2019-nCoV and naming it SARS-CoV-2. Nat Microbiol 2020;5:536–44.

[2] Wu F, Zhao S, Yu B, Chen YM, Wang W, Song ZG, et al. A new coronavirus associated with human respiratory disease in China. Nature 2020;579:265–9.

[3] Zhang Z, Song S, Yu J, Zhao W, Xiao J, Bao Y. The Elements of Data Sharing. Genomics Proteomics Bioinformatics 2020;18:1–4.

[4] Shu Y, McCauley J. GISAID: Global initiative on sharing all influenza data - from vision to reality. Euro Surveill 2017;22:30494.

[5] O’Leary NA, Wright MW, Brister JR, Ciufo S, Haddad D, McVeigh R, et al. Reference sequence (RefSeq) database at NCBI: current status, taxonomic expansion, and functional annotation. Nucleic Acids Res 2016;44:D733–45.

[6] Zhao WM, Song SH, Chen ML, Zou D, Ma LN, Ma YK, et al. The 2019 novel coronavirus resource. Yi Chuan 2020;42:212–21. (in Chinese with an English abstract)

[7] Tang X, Wu C, Li X, Song Y, Yao X, Wu X, et al. On the origin and continuing evolution of SARS-CoV-2. National Science Review 2020;7:1012–23.

[8] Korber B, Fischer W, Gnanakaran S, Yoon H, Theiler J, Abfalterer W, et al. Spike mutation pipeline reveals the emergence of a more transmissible form of SARS-CoV-2. bioRxiv 2020; https://doi.org/10.1101/2020.04.29.069054.

[9] Bandelt HJ, Forster P, Rohl A. Median-joining networks for inferring intraspecific phylogenies. Mol Biol Evol 1999;16:37–48.

[10] Croucher NJ, Didelot X. The application of genomics to tracing bacterial pathogen transmission. Curr Opin Microbiol 2015;23:62–7.

[11] Mavian C, Pond SK, Marini S, Magalis BR, Vandamme AM, Dellicour S, et al. Sampling bias and incorrect rooting make phylogenetic network tracing of SARS-COV-2 infections unreliable. Proc Natl Acad Sci U S A 2020;117:12522–3.

[12] Eyre DW, Cule ML, Wilson DJ, Griffiths D, Vaughan A, O’Connor L, et al. Diverse sources of C. difficile infection identified on whole-genome sequencing. N Engl J Med 2013;369:1195–205.

[13] Rambaut A, Holmes EC, O’Toole A, Hill V, McCrone JT, Ruis C, et al. A dynamic nomenclature proposal for SARS-CoV-2 lineages to assist genomic epidemiology. Nat Microbiol 2020. https://doi.org/10.1038/s41564-020-0770-5

[14] Kim D, Lee JY, Yang JS, Kim JW, Kim VN, Chang H. The Architecture of SARS-CoV-2 Transcriptome. Cell 2020;181:914–21 e10.

[15] Elbe S, Buckland-Merrett G. Data, disease and diplomacy: GISAID’s innovative contribution to global health. Glob Chall 2017;1:33–46.

[16] National Genomics Data Center M, Partners. Database Resources of the National Genomics Data Center in 2020. Nucleic Acids Res 2020;48:D24–D33.

[17] Shi W, Qi H, Sun Q, Fan G, Liu S, Wang J, et al. gcMeta: a Global Catalogue of Metagenomics platform to support the archiving, standardization and analysis of microbiome data. Nucleic Acids Res 2019;47:D637–D48.

[18] Xiao SZ, Armit C, Edmunds S, Goodman L, Li P, Tuli MA, et al. Increased interactivity and improvements to the GigaScience database, GigaDB. Database (Oxford) 2019;2019:1–9.

[19] Edgar RC. MUSCLE: multiple sequence alignment with high accuracy and high throughput. Nucleic Acids Res 2004;32:1792–7.

[20] McLaren W, Gil L, Hunt SE, Riat HS, Ritchie GR, Thormann A, et al. The Ensembl Variant Effect Predictor. Genome Biol 2016;17:122.

[21] Rego N, Koes D. 3Dmol.js: molecular visualization with WebGL. Bioinformatics 2015;31:1322–4.

