## Supplementary figures and images for "The global landscape of SARS-CoV-2 genomes, variants, and haplotypes in 2019nCoVR"

### Figure S1

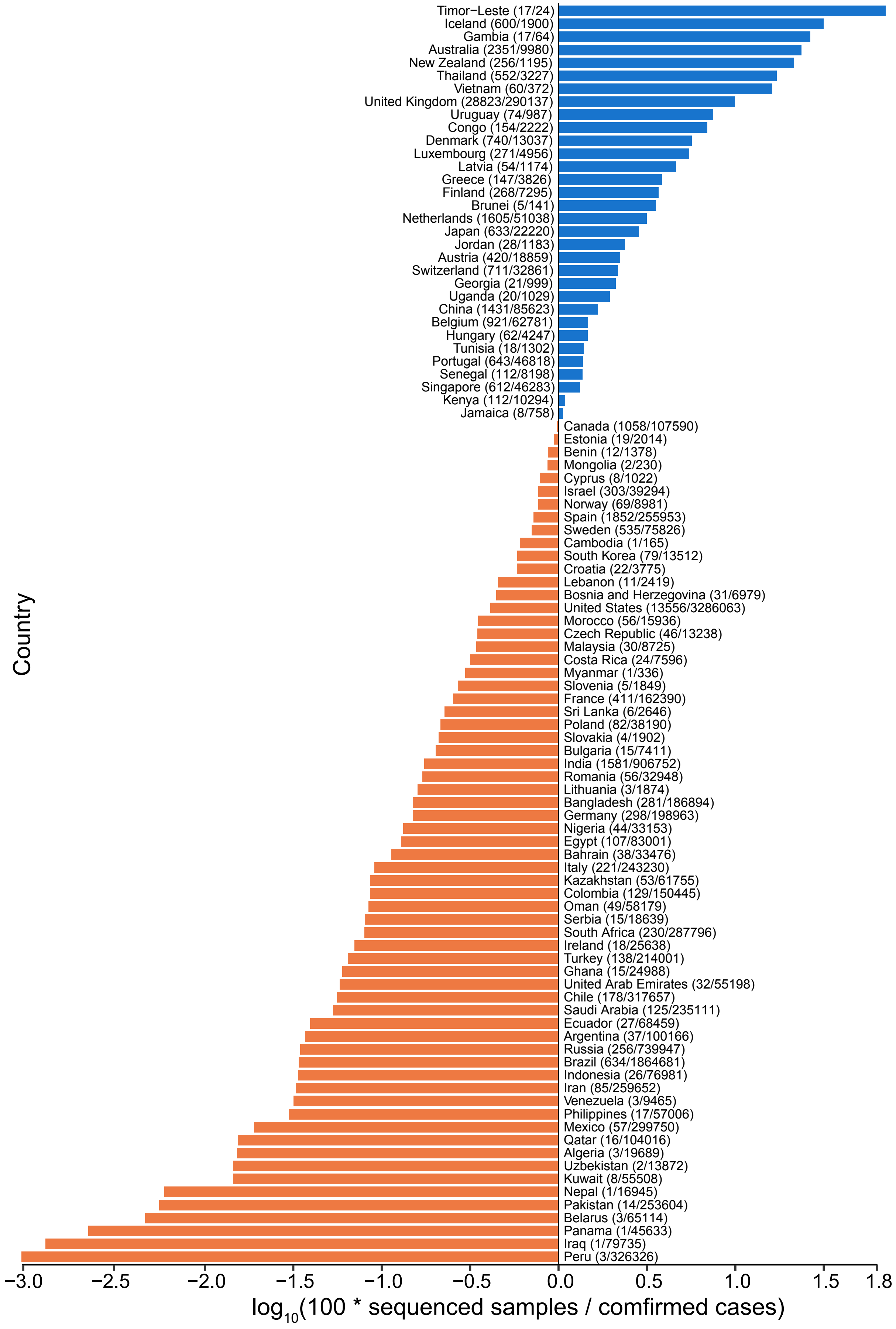

### Figure S2

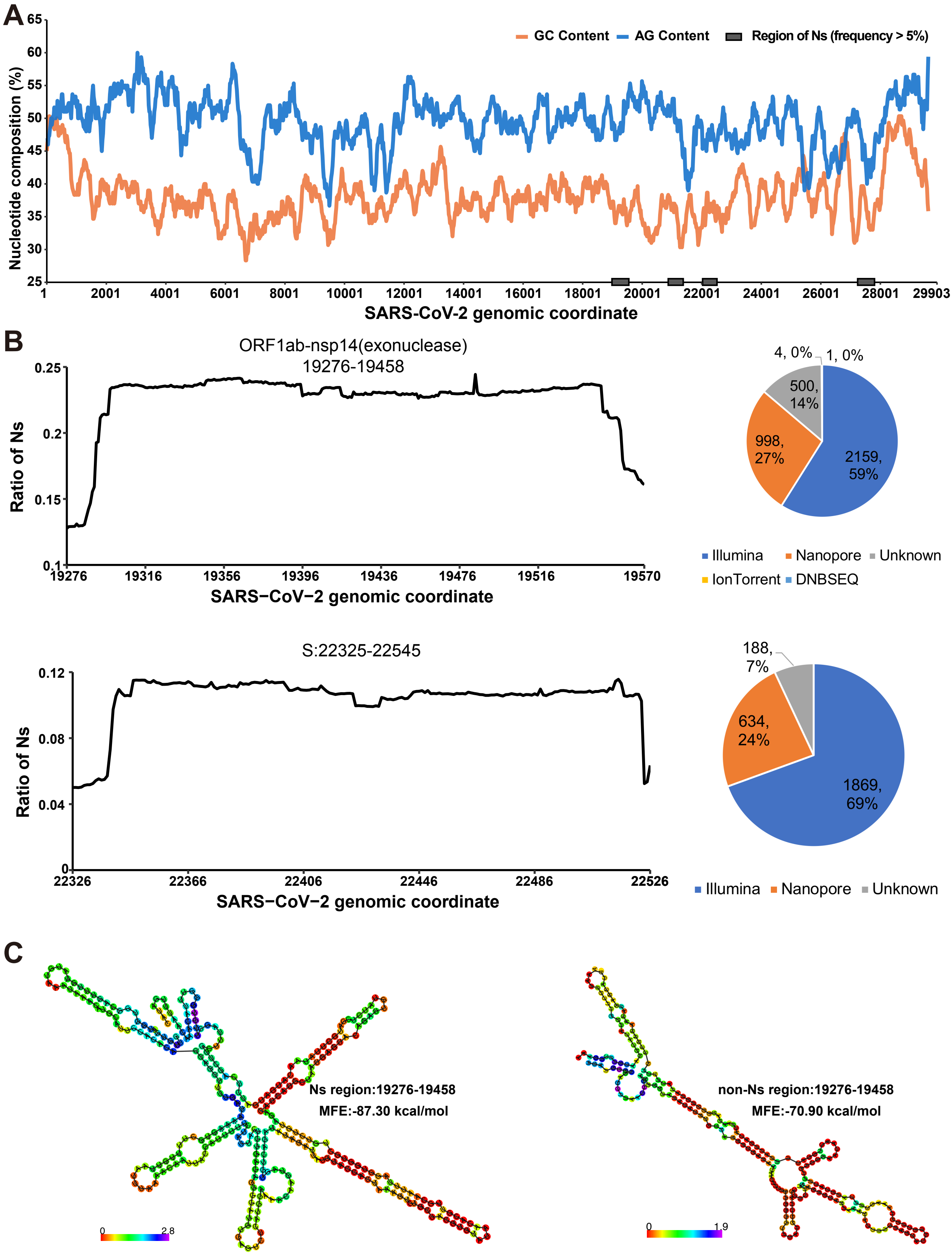
